# Glycemia Shift Pancreatic Islets Rhythmicity via δ-α Cell in vivo, Impairment in Diabetes

**DOI:** 10.1101/2025.08.09.669458

**Authors:** Yawen Deng, Zhenchao Fu, Xuejiao Wang, Yongxing Qiao, Xi Wu, Sen Yang, Chunmei Zhou, Wenlong Huang, Lijing Hui, Weiran Qian, Liangyi Chen, Chao Tang, Yuanyuan Du, Xiaohong Peng, Huixia Ren

**Affiliations:** Capital Medical University, Beijing 100069, China; Institute for Medical Physiology, Chinese Institutes for Medical Research (CIMR), Beijing, 100069, China; Hangzhou Institute of Medicine (HIM), Chinese Academy of Sciences, Hangzhou, Zhejiang, 310000, China; State Key Laboratory of Membrane Biology, Beijing Key Laboratory of Cardiometabolic Molecular Medicine, Institute of Molecular Medicine, School of Future Technology, Center for Life Sciences, Peking University, Beijing 100871, China; Center for Quantitative Biology, Peking University, Beijing 100871, China; Shenzhen University, Shenzhen 511464, China

**Author notes:** These authors contributed equally (Yawen Deng, Zhenchao Fu, Xuejiao Wang, Yongxing Qiao, Xi Wu).

**Keywords:** islet, blood glucose, Ca^2+^ oscillation, diabetes, Glp1r

## Abstract

Blood glucose homeostasis relies on the well-coordinated rhythmic activity of millions of islets throughout the pancreas. Islet rhythmicity is triggered by glucose elevation and mediated by paracrine interactions. However, the dynamics of islet population rhythmicity in healthy and diabetic pancreases in vivo remain poorly understood. Using simultaneous multi-islet Ca^2+^ imaging (20-100 islets per experiment) in both live mice and pancreatic tissue slices, we systematically studied how glycemia fluctuations and intra-islet paracrine signaling collectively shape the islet rhythmicity. In this study, we report that a transition from Hyperglycemia to Euglycemia induces a coordinated shift from slow to fast islet Ca^2+^ oscillations (HESF) in vivo. HESF is conserved in pancreatic tissue slices and isolated islets, however, not dispersed single cells in vitro, suggesting a mechanistic link with paracrine interactions. We found HESF arises from α-cell activation, which is inhibited by δ cells upon glucose elevation. The autonomous islets mostly differ in phase and period at high glucose level. Diabetic mice with disrupted glycemic stability lost HESF both in vivo and in vitro. Interestingly, HESF is preserved in β-cell knockout Gcgr transgenetic mice, both in vivo and in vitro, suggesting HESF’s dependence on Glp1r. Indeed, HESF was restored in semaglutide-treated diabetic mice with stabilized glycemic stability. These findings offer a comprehensive understanding of how δ and α cells influence islet rhythmicity and precisely maintain the stability of blood glucose.

## Introduction

Maintenance of blood glucose homeostasis is a fundamental physiological process, with pancreatic islets serving as critical functional units orchestrating this balance. The human pancreas harbors ∼1 million islets, while mice possess ∼1,000, each functioning as micro-organs composed of β cells (insulin-secreting), α cells (glucagon-secreting), and δ cells (somatostatin-secreting)^1^. Pancreatic islets function as a distributed network at the population level to stable blood glucose, relying on inter-islet coordination^2^ and intra-islet paracrine signaling^2–4^. The seminal observation that transplanting islets from different donor species drives recipients to adopt a glycemic set point matching that of the donors^5^ underscores the decisive role of islets in glucose regulation. Emerging evidence highlights the critical roles of α-cell^6^ and δ-cell^7^ in establishing the glycemic set point through paracrine signaling.

Beta-cells have a specialized mechanism to detect changes in blood glucose concentration^8,9^. This mechanism involves glucose entering the cell, being metabolized, and the elevated ATP/ADP shaping the cell’s oscillatory electrical activity and insulin secretion^10^. Islet rhythmicity^11–13^—oscillatory responses to glucose with periods ranging from 20 to 600 seconds—is central to glucose control, and disruptions in this rhythmicity are linked to diabetes pathogenesis^14–16^. While paracrine signaling is known to tune islet oscillatory patterns^17^, the spatiotemporal coordination of glucose-driven islet response—particularly the transition during glycemic shifts—remain uncharacterized. Moreover, the disruptions in islet rhythmicity lacks in vivo validation.

Most prior studies have focused on single-cell or single-islet dynamics^18^, leaving the population-level coordination of islets in vivo poorly understood. The major challenge in functional studies of the islet population in health and disease is the fact that they are distributed across the pancreas, which requires large focus of view and trans-genetic labeling.

For this purpose, we generated β-cell trans-genetic label GCaMP6f obese mutant mouse (*ob/ob*) as a model system. Using large field simultaneous multi-islet Ca²⁺ imaging (20–100 islets per experiment) in live mice and pancreatic slices, we uncover a hyperglycemia-to-euglycemia-induced shift from slow to fast islet Ca²⁺ oscillations (HESF). Mechanistically, HESF relies on α-cell activation, which is inhibited by δ cells during hyperglycemia. In diabetic mice with glycemic instability, HESF is disrupted both in vivo and in vitro. Strikingly, HESF is preserved in β-cell-deficient *Gcgr* transgenic mice and rescued by GLP-1 receptor agonist (semaglutide) treatment diabetic mice, linking this phenomenon to GLP-1 signaling. These findings establish a dynamic framework for how α-δ cell interactions encode glycemic information into islet rhythmicity, providing mechanistic insights into glucose homeostasis and its breakdown in diabetes.

## Results

### Hyperglycemia-to-Euglycemia Induces a Coordinated Slow-to-Fast Ca^2+^ Oscillations Transition in Pancreatic Islets in Vivo

To investigate how pancreatic islets coordinate rhythmic activity in vivo to maintain glycemic homeostasis, we developed a large-field imaging modality capable of observing 20–100 pancreatic islets within a 3.5×6.0 mm field of view (2x magnification) for over four hours (Fig. 1a). Intravenous glucose infusion was used to perturb glycemia and assess islet responsiveness, while intravenous glucose tolerance tests (IVGTT, 0.6 g/kg) evaluated glucose handing capacity.

To characterize glycemic homeostasis, mice with β cell specific GCaMP6f expression (*Ins1-Cre^+/-^; GCaMP6f^f/+^*) were implanted with continuous glucose monitoring (CGM) devices. CGM data revealed euglycemia ranges of 4-7 mM, with fluctuations of 0.9–1.5 mM (Fig. 1b). The mean glucose level detected by CGM nicely correlated with tail-vein fasting glucose level in mice (Fig. 1c), validating the reliability of CGM for tracking in vivo glycemia.

In vivo, islets exhibited robust rhythmic Ca^2+^ activity, with oscillation modes dynamically regulated by blood glucose levels (Fig.1d). At euglycemia level, 58% of islets displayed fast Ca^2+^ oscillations and remaining as slow. During IVGTT, all islets responded to acute glycemic elevation: 77% with sustained Ca²⁺ elevation and 23% with slower Ca^2+^ oscillations in the first 30 min, then all islets showed slow oscillations. The peak blood glucose level during IVGTT correlated with each mouse’s euglycemia level, confirming interindividual differences in glucose tolerance (Fig. S1a).

Following islets activation, blood glucose subsequently declined. This transition from Hyperglycemia to Euglycemia triggered a coordinated Slow-to-Fast Ca^2+^ oscillation transition (termed HESF), with 65% of islets stabilizing in a fast-oscillation state (Figs. 1d, S1b and Video 1). Notably, the glucose levels trigging fast oscillations strongly correlated with CGM-detected euglycemia levels (Fig. 1e), suggesting that HESF is tightly linked to physiological glycemic set points. Mice with chronically elevated baseline glycemia exhibited a higher glucose threshold for this transition, and vice versa (Figs. 1e and S1b). The amplitudes of the fast Ca²⁺ oscillations were significantly lower than that of the slow Ca²⁺ ones (Fig. S1b). Glucose clamping at 7G, 10G and 20G significantly increased oscillation period (Figs. 2a and 2b).

**Fig.1.**
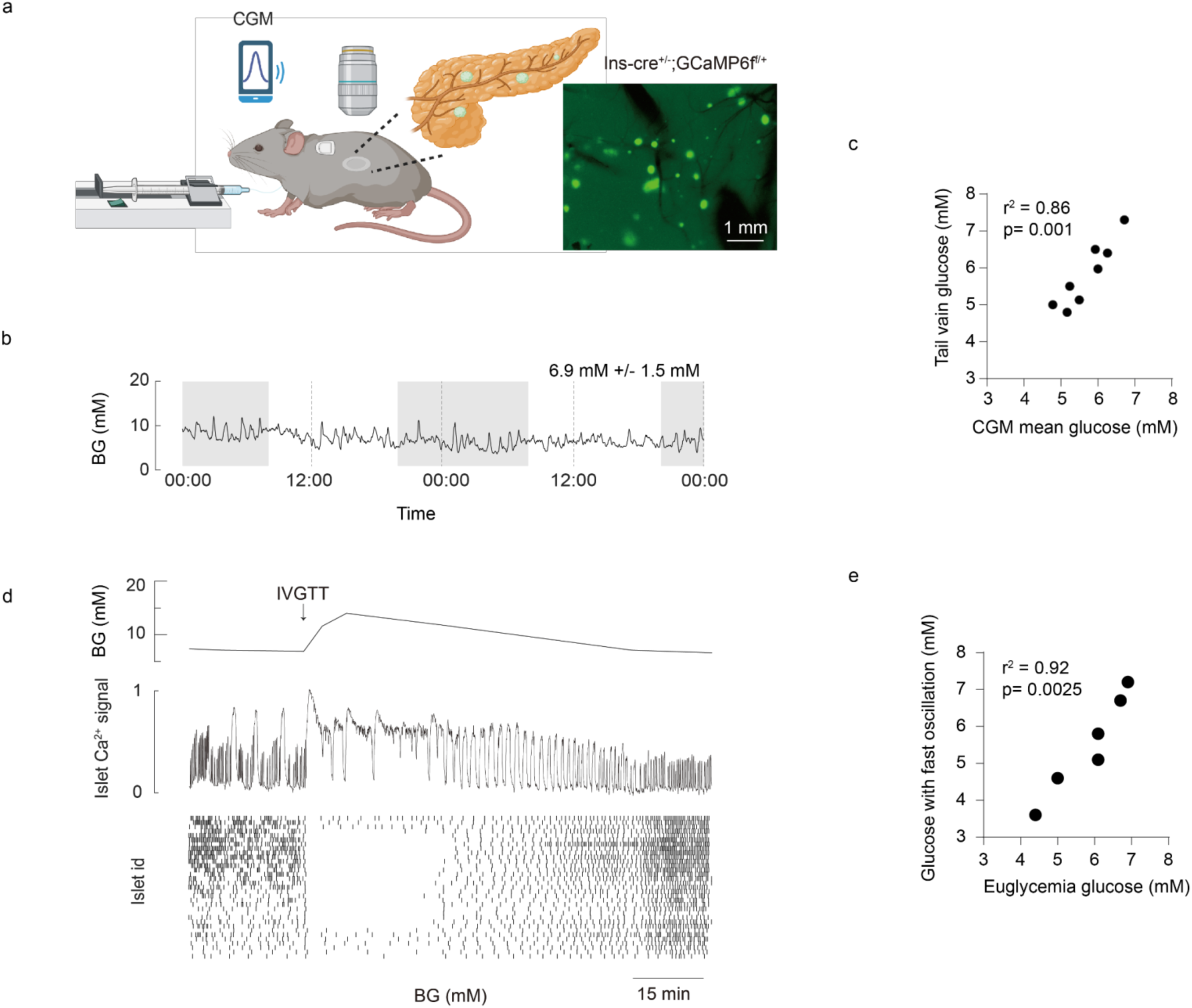
Hyperglycemia-to-Euglycemia Induces a coordinated Slow-to-Fast (HESF) Islet Ca^2+^ Oscillation Transition In Vivo. a) Experimental setup and islet imaging. Left: Schematic of in vivo Ca^2+^ imaging of pancreatic islets during hyperglycemic clamp. A continues glucose monitoring (CGM) device was implanted to measure glycemia levels in mice. Right: Maximum intensity projection of pancreatic islets from *Ins1-Cre^+/-^; GCaMP6f^+^* mice. b) Blood glucose profile. 48-hour CGM traces from one representative mice (dashed lines: 00:00/12:00; grey shading: nighttime 20:00–8:00). The number indicates the mean ± standard deviation of the representative CGM profile. c) Glucose measurements comparing CGM-derived mean values with tail vein blood glucose. d) Top: Blood glucose concentration. Glycemic challenges: resting phase and intravenous glucose tolerance test (IVGTT) phase. Middle: Representative islet Ca^2+^ trace. Bottom: Raster plot of oscillation peaks in 34 simultaneously monitored islets. e) Euglycemia (mean 48-hour CGM shown in b) vs. glucose with fast oscillation (mean glucose for periods ≤60 s shown in d); n = 6 mice. Yellow dot denotes mice in b and d.

Mostly, each islet functioned as an autonomous oscillator, differing in phase and period (Fig. S1c). Collectively, these data demonstrate that glycemic fluctuations dynamically tune the oscillation period of the islet population – with hyperglycemia inducing slow oscillations and euglycemia promoting fast oscillations.

### HESF is conserved in Pancreas Tissue Slices in Vitro and Dependent on Paracrine Signaling

Islets integrate local (paracrine) and systemic (hormonal/neural) signals to maintain glucose homeostasis. To elucidate the mechanisms underlying HESF, we next investigated islet Ca^2+^ activity in pancreas tissue slices that kept an intact surrounding without blood circulation and neuromodulation (Fig. 2a and S2a). As shown in Fig. 2a, islets in tissue slices showed stable basal Ca^2+^ levels in 5 mM glucose. Glucose elevation to 7 mM induced regular Ca^2+^ oscillations in islets (Fig. S2b). Notably, in islets with oscillations further glucose elevation to 10 and 20 mM markedly prolonged the Ca^2+^ oscillation period (Fig. 2c) like in vivo (Fig. 2b). Glucose elevation increased the Ca^2+^ amplitude (Fig. S2d). Exocrine cells from mice expressing the calcium indicator GCaMP6f also displayed repetitive Ca^2+^ signaling with much shorter oscillation periods, but unlike in the islets, the period was independent of the glucose concentration (Fig. S2e). We next investigated isolated islets, which retain paracrine interactions. The glucose modulated oscillation frequency is preserved, mirroring the responses of islets in tissue slices and in vivo (Fig. 2a and 2d). In contrast, dispersed single cells - lacking paracrine communication - lost HESF (Figs. 2a and 2e). Collectively, these data demonstrate that glucose dependent tuning of islet oscillation frequency is conserved across in vivo and in vitro models, provided that paracrine interactions remain intact.

**Fig. 2.**
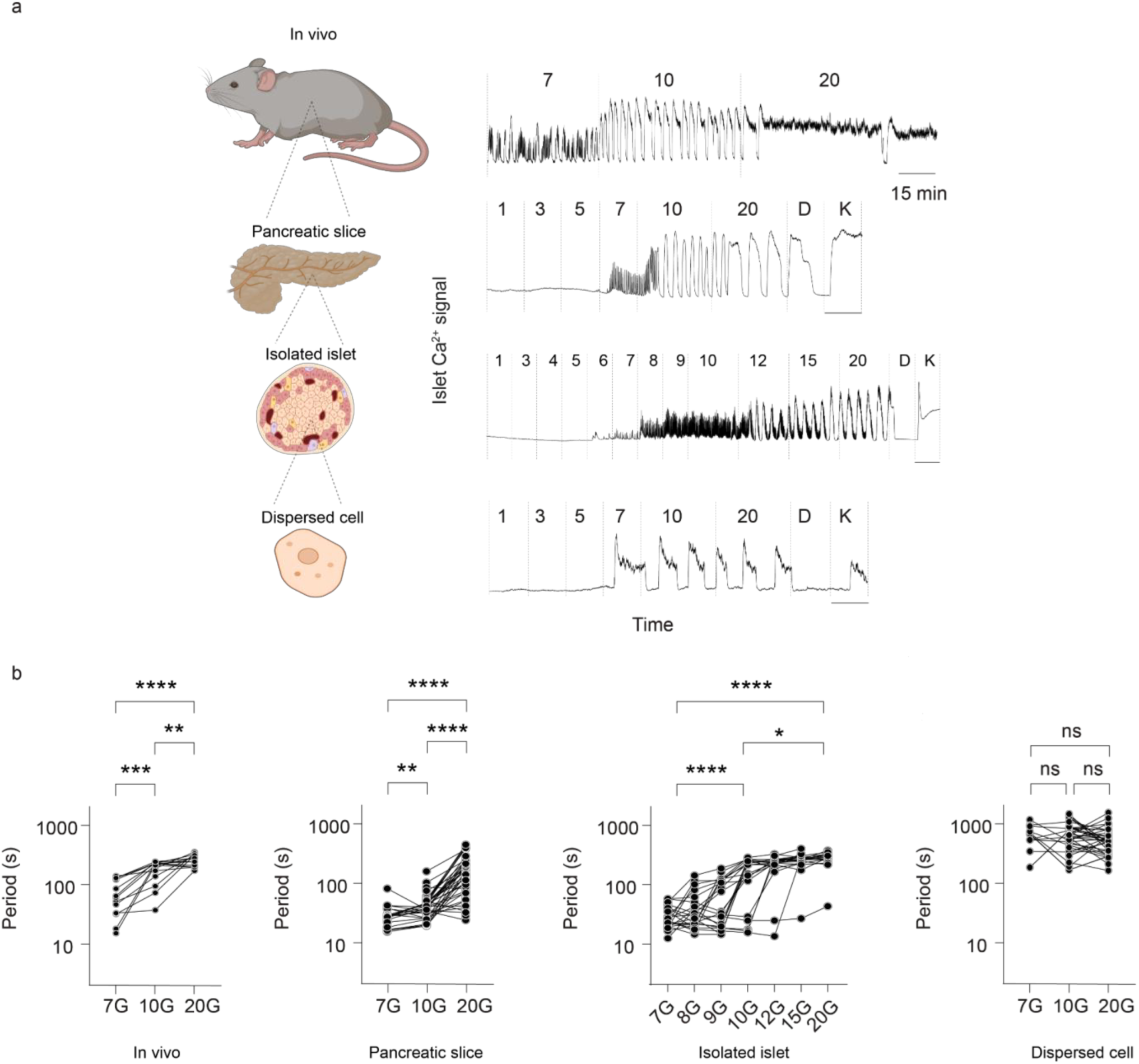
Conservation of HESF in pancreatic tissue slice, not in dispersed cell in Vitro. a) Schematic (left) and representative islet Ca^2+^ recordings (right) in vivo (top), pancreatic slice (middle) and dispersed cell (bottom) from *Ins1-Cre^+/-^; GCaMP6f^+^* mice. The sequential perfusion includes low glucose (1G, 3G; 15 min each; not shown), physiological glucose (5G, 7G; 15 min each), high glucose (10G, 20G; 30 min each), and controls (25 mM KCl, 250 uM diazoxide). b) In vivo islets Ca^2+^ oscillation periods at 7G, 10G and 20G (n = 93 islets from 3 mice). c) Pancreatic slice Ca^2+^ oscillation periods at 7G, 10G and 20G (n = 20, 44, 83 islets from 9 mice). d) Isolated islet Ca^2+^ oscillation periods at 7G, 8G, 9G, 10G, 12G, 15G and 20G (n = 22 islets from 4 mice). e) Dispersed β cell Ca^2+^ oscillation periods at 7G, 10G and 20G (n = 50 cells from 3 mice).

### Islet Rhythmicity is modulated by δ-α cell interactions

The conservation of HESF in pancreas tissue slice and isolated islet suggests the glucose-modulated oscillation modes were caused by intra-islet paracrine communication. We found that somatostatin prolonged the oscillation cycles from 23 s to 500 s at 10G (Fig. 3a), whereas glucagon instead reduced the oscillations periods at 20G from 73 s to 31 s (Fig. 3b). These findings indicate that paracrine influence from glucagon and somatostatin modulate the transition between slow and fast oscillations. Indeed, blocking the somatostatin receptors 2 on the glucagon-releasing α-cell with CYN154806 resulted in faster oscillations at 20G (cycles reduced from 79 to 50 s Fig. 3C), indicating that inhibition of glucagon secretion by endogenous somatostatin was relieved by the receptor antagonist.

**Fig. 3.**
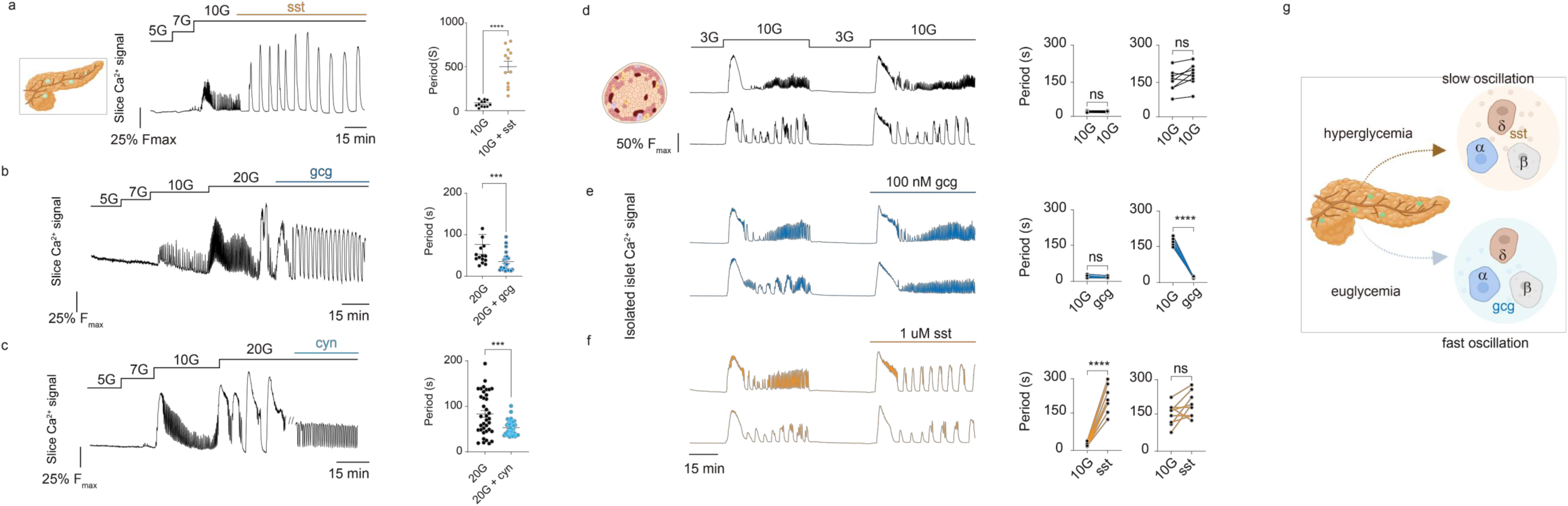
HESF arising from α and δ cell paracrine interactions. a) Left: Ca^2+^ trace in pancreatic tissue slice under a stimulation protocol with 5G, 7G, 10G and 1 uM somatostatin. Right: Ca^2+^ oscillation periods at 10G and somatostatin (n = 12 islets from 3 mice). b) Left: Ca^2+^ trace in pancreatic tissue slice under a stimulation protocol with 5G, 7G, 10G, 20G and 1 uM glucagon. Right: Ca^2+^ oscillation periods at 20G and glucagon (n = 23 islets from 3 mice). c) Left: Ca^2+^ trace in pancreatic tissue slice under a stimulation protocol with 5G, 7G, 10G, 20G and 10 uM cyn (CYN 154806). Right: Ca^2+^ oscillation periods at 20G and cyn (n = 37 islets from 3 mice). d) Isolated fast and slow islet Ca^2+^ signal with 3G, 10G, 3G and 10G perfusion (left). Period changes in fast (middle) vs. slow (right) islets during sequential 10G stimulation (n = 9 fast and 8 slow islets from 3 mice). e) Same as d), 100 nM glucagon added during second 10G period (n = 7 fast and 10 slow islets from 3 mice). f) Same as d), 1 uM somatostatin added during second 10G period (n = 7 fast and 9 slow islets from 3 mice). g) Proposed HESF mechanism. Euglycemia → α-cell glucagon secretion → fast oscillation. Hyperglycemia → δ-cell somatostatin secretion → slow oscillation.

Also isolated islets showed characteristic individual Ca^2+^ patterns. Repetitive 10G stimulation thus induced different but consistent oscillation modes in individual islets (Fig.3d). Glucagon accelerated slow islet oscillations into fast ones without affecting already fast oscillations (Fig. 3e). Conversely, somatostatin transformed fast islet oscillations into slow ones while leaving slow-oscillating islets unaltered (Fig. 3f). These observations indicated that δ and α cells form the core regulatory network for the intrinsic rhythmicity of the islet (Fig. 4g).

**Fig. 4.**
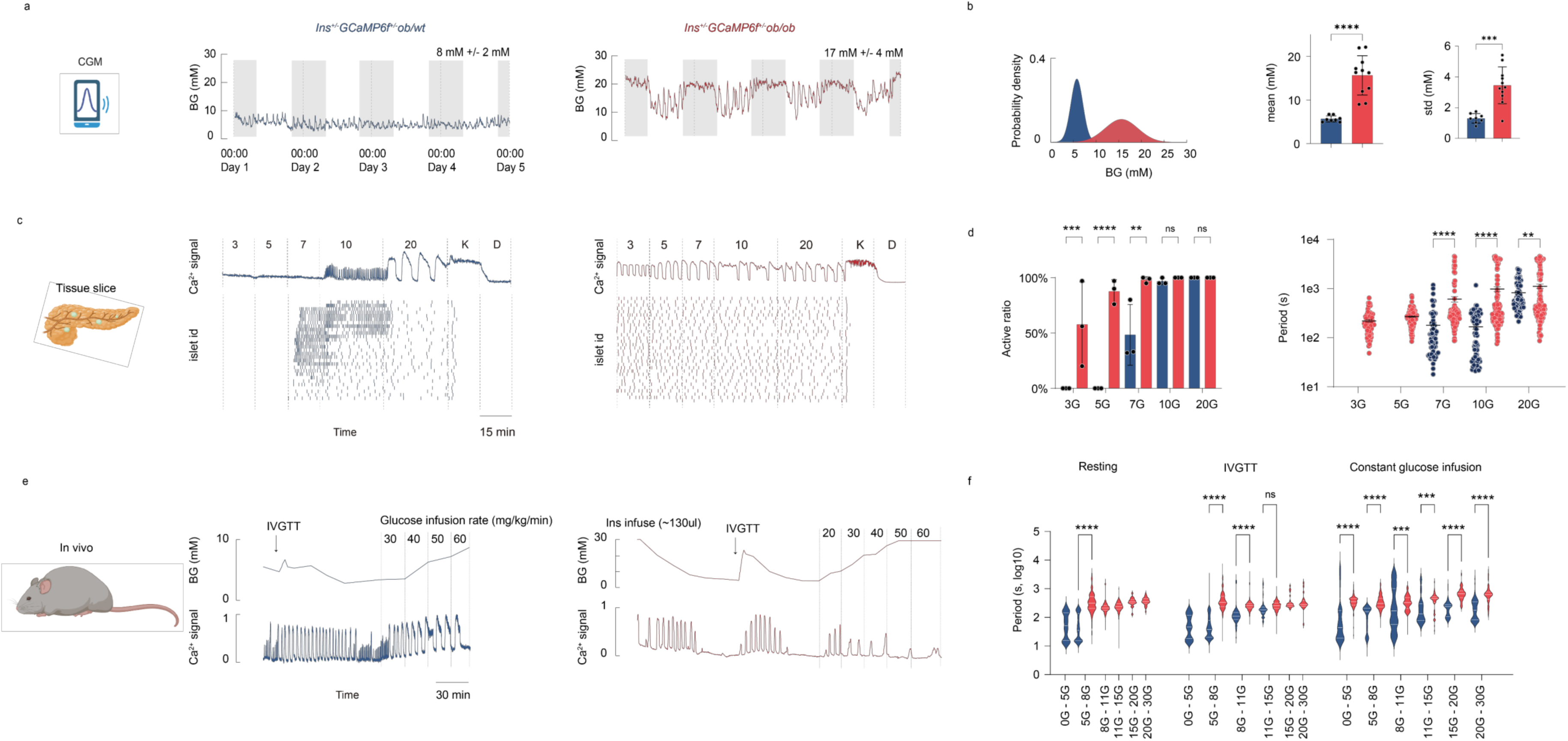
Loss of Glycemic Stability and HESF in Diabetic Mice. a) Four-day CGM from control (top, *Ins1^+/-^;GCaMP6f^f/+^;ob/wt*, blue) and diabetic mice (bottom, *Ins1^+/-^;GCaMP6f^f/+^;ob/ob*, red). Grey shading: nighttime 20:00–8:00. Number shows mean ± standard derivation. b) Blood glucose distribution profiles from a) (left). Quantitative CGM analysis comparing mean glucose (middle), standard derivation (right), n = 8 control mice and 11 diabetic mice. c) Pancreatic tissue slice imaging of control (left) and diabetic (right) mice. Characteristic islet Ca^2+^ traces and raster plots of oscillation peaks during sequential perfusion of 3G, 5G, 7G, 10G and 20G, followed by KCl and diazoxide. d) Active ratio and oscillation period comparison for islets in c) (n = 93 islets from 3 *Ins1^+/-^;GCaMP6f^f/+^;ob/wt* mice, n = 154 islets from 3 *Ins1^+/-^;GCaMP6f^f/+^;ob/ob* mice). e) In vivo imaging of control (left) and insulin-infused diabetic mice (right). Blood glucose and islet Ca^2+^ trace during resting phase, IVGTT phase and constant glucose infusion phase. f) Oscillation period comparison for islets in e) (n = 84 islets from 3 *Ins1^+/-^;GCaMP6f^f/+^;ob/wt* mice and n = 73 islets from 3 insulin-infused *Ins1^+/-^;GCaMP6f^f/+^;ob/ob* mice).

### Diabetic mice exhibit elevated glycemic fluctuations

We next characterized the glycemic profiles of diabetic *ob/ob* mice and their normal control *ob/wt* littermates (Fig. 4a). Control *ob/wt* mice exhibited stable glycemia levels with limited variations (Fig. 4b, 8 ± 2 mM). In contrast, *ob/ob* mice showed chronically elevated glycemia accompanied with substantial fluctuations (Fig. 4b, 17 ± 4 mM). Notably, during the normal nocturnal feeding period *ob/ob* mice failed to regulate glycemia and instead sustained persistently elevated levels (∼ 20 mM). These observations reveal that diabetic *ob/ob* mice have a diminished capacity to counteract feeding-induced glycemic perturbations, highlighting a critical defect in their glycemic homeostatic machinery.

### Impaired islet function in diabetic mice: abolished fast Ca²⁺ oscillations and blunted glucose sensitivity

Islets from *ob/ob* mice pancreatic slice exhibited abnormal activation patterns (Figs. 4c and 4d). Unlike in the controls they showed Ca^2+^ oscillations at significantly lower glucose concentrations (Fig. 4d). We quantified the ratio of active, fast (period <1 min), slow (period 1-15 min) and sustained (period >15 min) Ca^2+^ patterns (Fig. 4d). At 3 mM and 5 mM glucose, all *ob/wt* islets were inactive (Fig.4d). 58% *ob/ob* islets were active at 3 mM glucose and increased to 88% at 5 mM glucose. At 7 mM glucose, 48% *ob/wt* islets were active (20% showed fast oscillations and 28% slow). Nearly all *ob/wt* islets (97%) response to 10G with fast (38%) or slow (58%) oscillations. In 20G most *ob/wt* islets (68%) showed slow oscillations and 32% more sustained Ca^2+^ patterns. However, only slow oscillations were observed in *ob/ob* islets irrespective of glucose concentration. In 10-20 mM glucose the slow oscillations were transformed into Ca²⁺ elevation in 27% of the *ob/ob* islets.

Ca^2+^ dysregulation was also apparent in *in vivo* imaging experiments (Fig. 4e). Elevating blood glucose by an IVGTT failed to induce significant changes in *ob/ob* islet Ca²⁺ dynamics or population activity (Fig. S3).

To further characterize how *ob/ob* islets respond under glycemic levels comparable to those of *ob/wt* mice, we performed reciprocal glycemic manipulations: exogenous glucose infusion elevated blood glucose in *ob/wt* mice to hyperglycemic levels (Fig. 4e, left panel), while insulin administration lowered blood glucose in *ob/ob* mice to the normoglycemic range (Fig. 4e, right panel).

At resting phase, *ob/wt* islets exhibited Ca²⁺ oscillations under euglycemia, a feature that was completely absent in *ob/ob* islets (Fig. 4f, Resting phase). Across a broad glycemic range (5 – 30 mM), *ob/ob* islets displayed glucose-insensitive oscillation period (∼300 s), in stark contrast to the glucose-dependent tuning observed in controls. As blood glucose levels decreased, slow Ca²⁺ oscillations in *ob/ob* islets became inactive.

A subsequent intravenous glucose bolus (0.6 g/kg) increased glucose to 9 mM in *ob/wt* mice and 15 mM in *ob/ob* mice, triggering robust slow Ca²⁺ oscillations in both groups. Over the following ∼1 hour, as blood glucose declined, *ob/wt* islets transitioned back to fast oscillations, whereas *ob/ob* islets maintained significantly slower oscillations before reverting to an inactive state. This confirms that HESF is lost in *ob/ob* mice in vivo.

During the constant glucose infusion phase, 40% of *ob/wt* islets exhibited slower oscillations, while 60% showed sustained Ca²⁺ elevation. In contrast, *ob/ob* islets remained insensitive to glucose stimulation (Figs. 5m and 5n).

Collectively, *ob/ob* islets displayed two key abnormalities compared to controls: complete loss of fast Ca²⁺ oscillations at euglycemic levels, and development of glucose insensitivity during hyperglycemia thus loss HESF, observed consistently both in vivo and in vitro.

**Fig.5.**
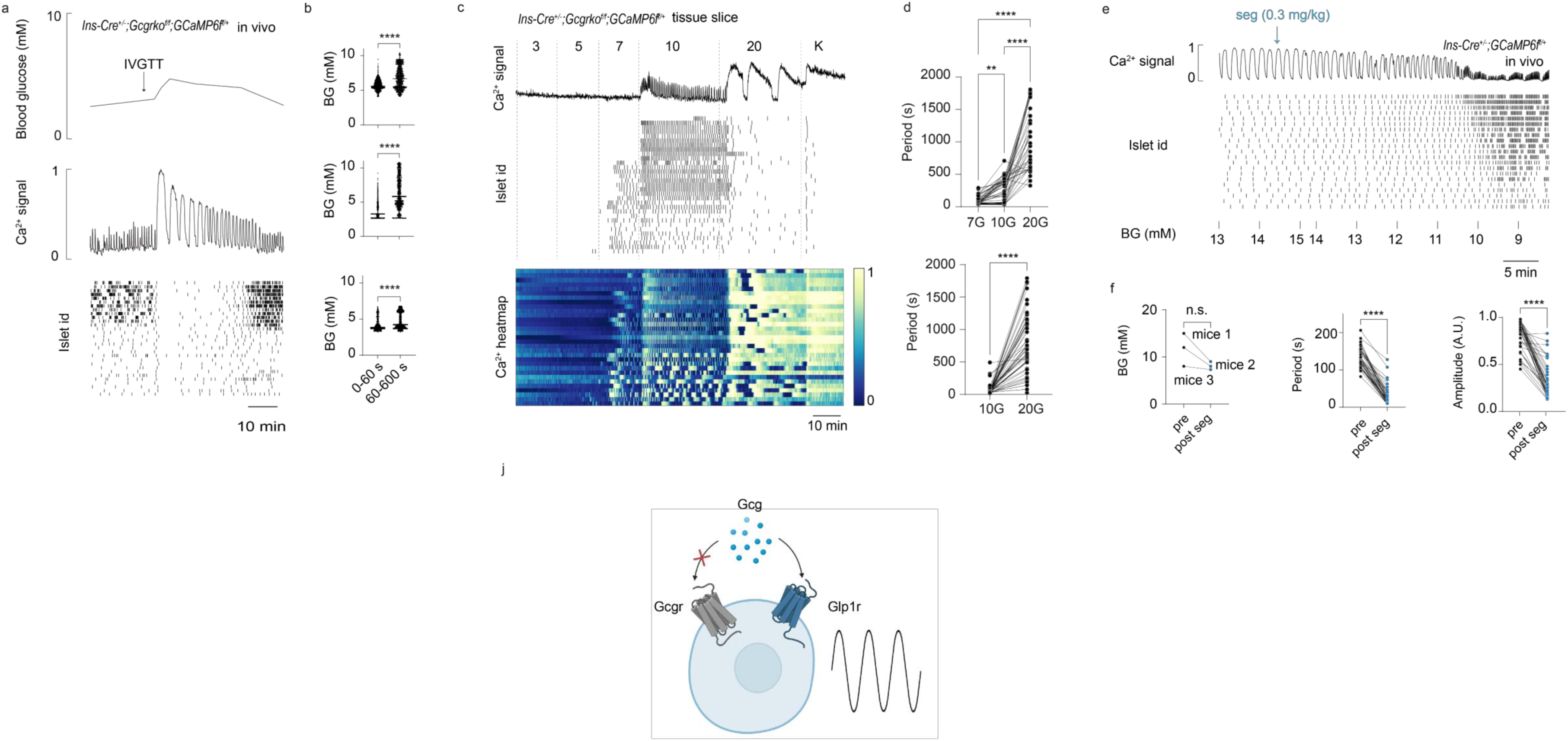
Fast islet Ca^2+^ Oscillation arises from endogenous activation of GLP1r. a) Blood glucose (top), in vivo islet Ca^2+^ signal (middle) and raster plots of Ca^2+^ oscillation peaks (bottom) in *Ins1-Cre^+/-^;Gcgrko^f/f^;Gcamp6f^f/+^* mice during resting and IVGTT phase. b) Blood glucose for islets exhibiting fast and slow oscillations. c) Representative islet Ca^2+^ trace, raster plots and heatmap in *Ins1^+/-^; Gcgrko^f/f^;GCaMP6f^f/+^* tissue slice during sequential perfusion of 3G, 5G, 7G, 10G and 20G, followed by KCl. d) Ca^2+^ oscillation period of 7G-responsive islets under 7G, 10G and 20G stimulation (left, n = 68 islets from 3 mice) and 10G-responsive islets under 10G and 20G stimulation (right, n = 60 islets from 3 mice). e) Representative in vivo Ca^2+^ trace, raster plots and Ca^2+^ peaks and blood glucose pre/post semaglutide in *Ins1^+/-^;GCaMP6f^f/+^* mice. f) Quantified changes in blood glucose, mean oscillation period, Ca^2+^ amplitude pre/post semaglutide (n = 40 islets from 3 mice). g) Mechanistic model: Glucagon induces fast oscillations via Glp1r activation.

### Fast Ca^2+^ Oscillation Arise from endogenous glucagon Glp1r activation in islets

The glucagon effect on β-cells is mediated both by the receptors for glucagon (GCGR) and GLP-1 (GLP1R)^19–21^. To test if HESF is maintained in the absence of glucagon receptor, we analyzed islets Ca²⁺ dynamic in β cell specific knockout glucagon receptor (*Ins1-crer^+/-^;Gcgrko^-/-^;Gcamp6f^+/+^)* mice. Both in vivo (Figs. 6a-6c) and in vitro (Figs. 6d-6f), these mice retained HESF. Under basal conditions, 24% *Ins1-crer^+/-^;Gcgrko^-/-^;Gcamp6f^+/+^*islets showed stable fast Ca²⁺ oscillations. Following an IVGTT, islets showed slow oscillation, then shift to fast Ca²⁺ oscillations in 29% of islets (Fig. 6b). In pancreatic slices, 20 mM glucose markedly slowed oscillation in islets (Fig. 6e) but fast oscillations remained in 42% of islets exposed to 10 mM glucose (Fig. 6f). These findings indicate that endogenous GLP1R activation is sufficient to induce fast Ca²⁺ oscillations during euglycemia. Consistent with this, intraperitoneal GLP-1 agonist (0.3 mg/kg Semaglutide) induced stable fast Ca²⁺ oscillations in vivo (Figs. 6g-6i). Together, our results demonstrate that endogenous GLP1R activation triggers HESF (Fig. 6j).

**Fig.6.**
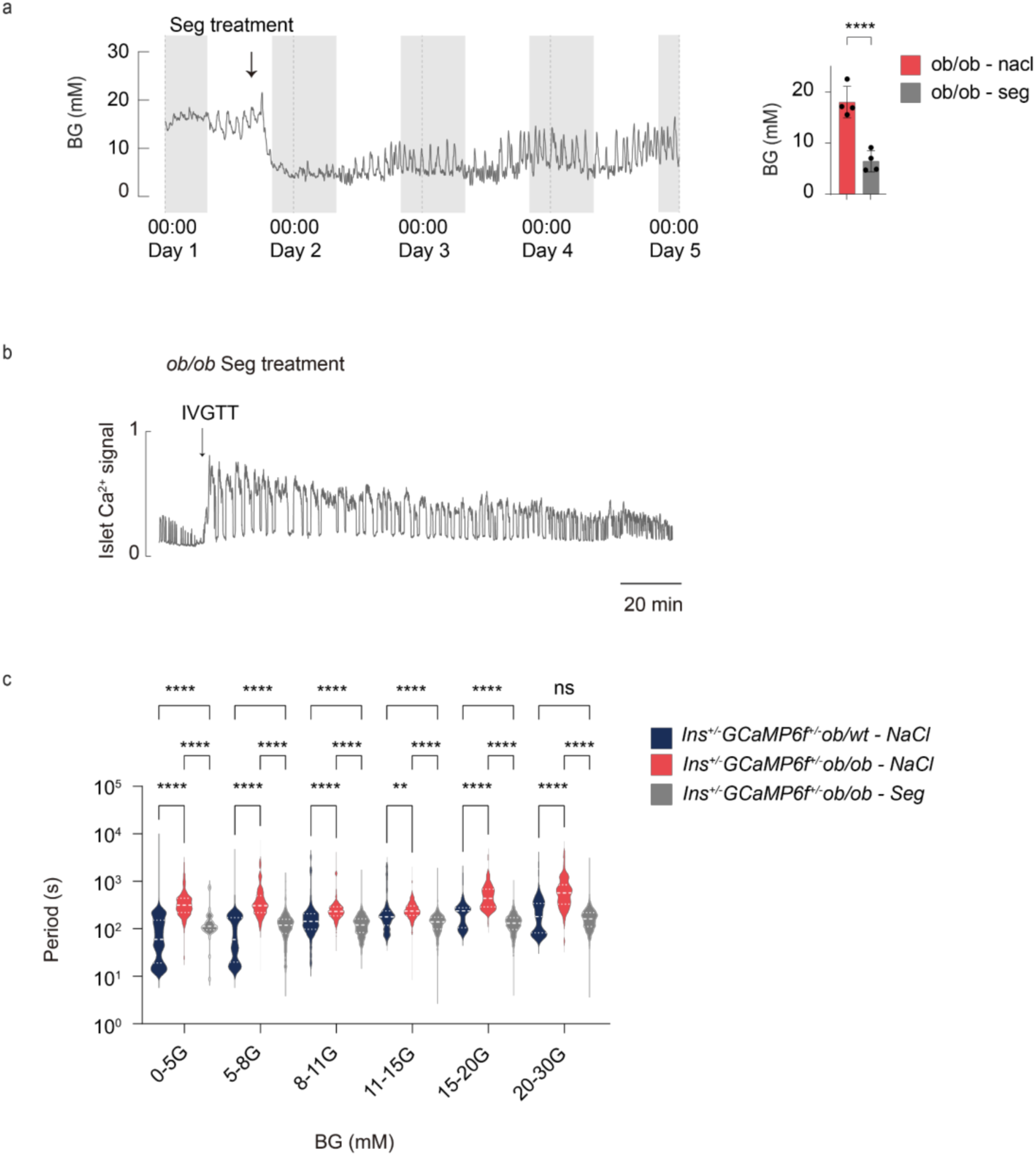
Glp1r treatment restores HESF. a) Left panel: Four-day CGM profiles from diabetic mice with semaglutide injection (*Ins1^+/-^ ;GCaMP6f^f/+^;ob/ob*, grey). Gray shading: nighttime 20:00–8:00; arrow: injection timepoint. Right panel: Mean blood glucose in semaglutide vs. saline injection diabetic mice. b) Representative in vivo Ca^2+^ trace from semaglutide treated *Ins1^+/-^;GCaMP6f^f/+^;ob/ob* mice. c) Islet oscillation periods in saline-treated control mice (n = 84 islets from 3 mice), saline-treated diabetic mice (n = 73 islets from 3 mice), and semaglutide-treated diabetic mice (n = 72 islets from 3 mice).

### HESF was restored in semaglutide-treated diabetic mice with stabilized glycemic stability

Given the impaired HESF in *ob/ob* mice, we investigated whether GLP1R signaling activation could restore fast Ca²⁺ oscillations and glycemic stability. Semaglutide injection rapidly reduced blood glucose from 18 mM to 6 mM within 1 hour (Figs. 6a-6d).

To evaluate longer-term effects, we administered GLP-1 agonist treatment to *ob/ob* mice for one week and assessed islet responses *in vivo*. We found at near-euglycemic glucose levels (5-8 mM) and islets Ca²⁺ oscillations with periods of 80-158 s (Figs. 6c) which is more similar to the periods in islet from control *ob/wt* mice (20–172 s), in contrast to the untreated *ob/ob* mice (219-498 s). During IVGTT-induced hyperglycemia the fast oscillations in GLP-1 agonist treated *ob/ob* mice were transform into slow ones but the faster pattern reappeared as the glucose levels normalized.

These findings indicate that GLP-1 treatment effectively restores normal euglycemia Ca²⁺ oscillation patterns with HESF function in the *ob/ob* mouse diabetes model.

### Fast Ca²⁺ Oscillations are Physiologically Stable

Finally, we studied the stability of islet’s fast oscillation (Fig. 7a). A stable state tends to return to its original state when slightly perturbed, whereas an unstable state tends to move away from its original state. Neutral state represents a passive state depends on perturbation level. We perturbed the islets by infusing pulses of glucose.

**Fig. 7.**
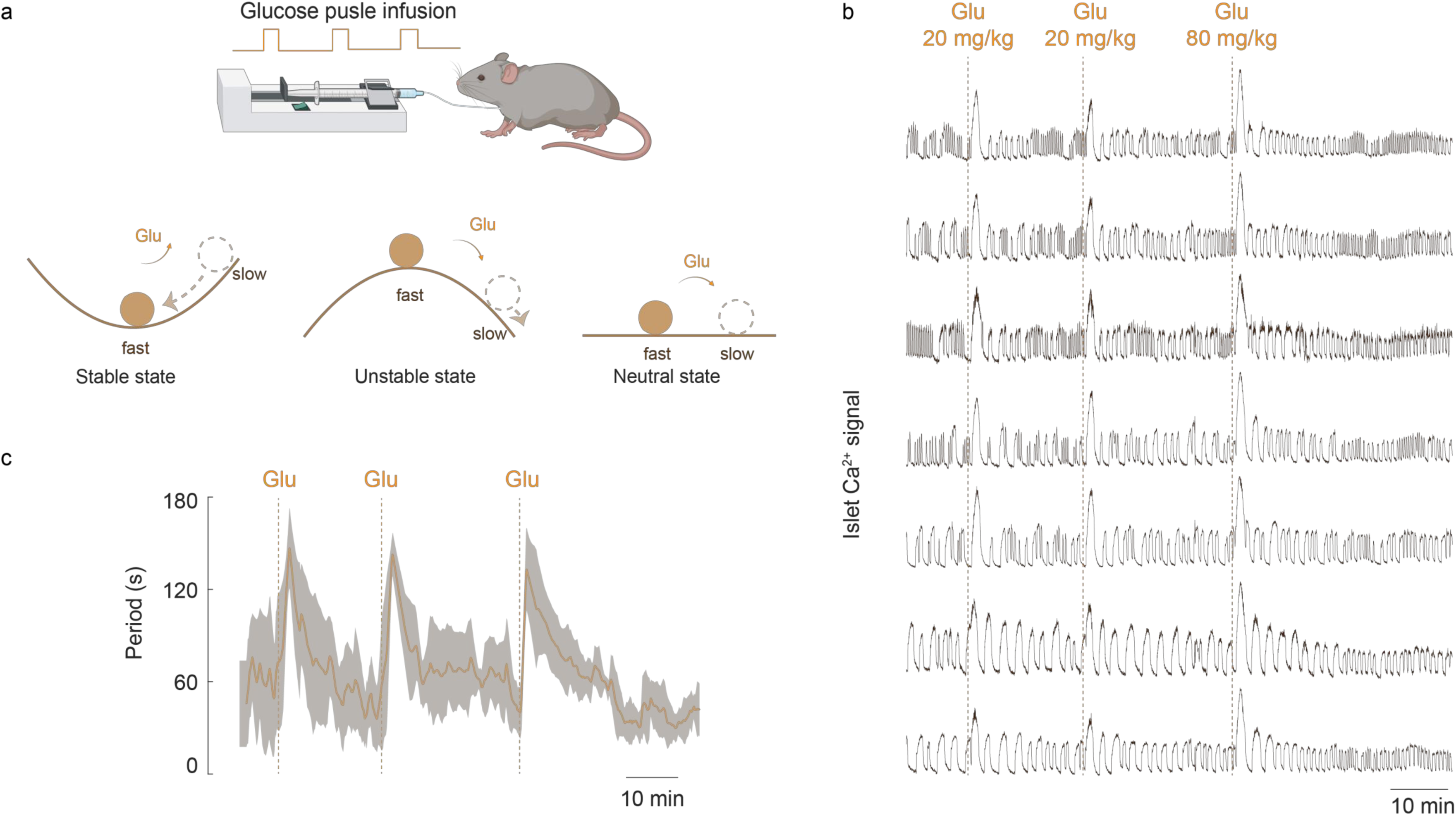
Fast Oscillations Represent a Physiologically Stable State in Vivo. a) Experimental design and stability concepts. Top: Schematic of glucose pulse perturbation protocol in vivo. Bottom: Theoretical response patterns for stable equilibrium (returns to initial state), unstable equilibrium (diverges from initial state) and neutral equilibrium (maintains perturbed state). b) Ca^2+^ traces from individual pancreatic islets before, during and after glucose pulse perturbations of islets with fast Ca^2+^ oscillations. c) Quantification of Ca^2+^ oscillation periods (means ± standard derivation) corresponding to traces in b.

As shown in Fig. 7b, repetitive infusions of 20 or 80 mg/kg glucose synchronously perturbated the fast oscillatory islets into slow ones with periods ∼150 s. Then the oscillation periods gradually shifted back to fast oscillations within ∼15 minutes (Fig. 7c). This suggests fast Ca²⁺ oscillations maintain a balanced glycemic influx and efflux at the set point, where slow Ca²⁺ oscillations and sustained Ca²⁺ elevation correspond to a larger efflux than influx, leading to a decrease in blood glucose. Fast oscillation is a physiological stable state.

## Discussion

Our study shows that blood glucose levels modulate the rhythmicity of pancreatic islet populations *in vivo*. Under euglycemic conditions, fast Ca²⁺ islet oscillations representing the beta cell activity are triggered by the activation of α-cells. In contrast, hyperglycemia initiates slow Ca²⁺ oscillations by engaging δ-cells and suppressing α-cell activity. Notably, in diabetic *ob/ob*-mice, where glycemic stability is compromised, the fast Ca²⁺ oscillations normally observed at euglycemia-induced were absent. Activation of Glp1r by semaglutide restored the fast Ca²⁺ oscillations in diabetic islets as well as glycemic stability in the diabetic mice.

Pancreatic tissue slices offer a experimental platform that bridges the gap between in vivo and in vitro isolated islet studies. Like *in vivo* islets in slices show fast and slow Ca²⁺ oscillations when stimulated by increasing glucose concentrations. In normal mice the fast oscillations dominated at the somewhat different glucose concentration that characterized euglycemia in each animal. Glucagon^22,23^ has long been known to promote fast islet Ca²⁺ oscillation, whereas somatostatin induces a slow pattern^17,24^. Our present studies extended the understanding of these interactions by relating the effects of glucagon and somatostatin to the prevailing glucose concentration.

Early *in vivo* studies of islet function by recording the membrane potential of beta cells in the pancreas confirmed earlier *in vitro* observations that an increase in glucose concentration extends the duration of the depolarized phase^25^. Subsequently, imaging methods have been developed to study islet activity *in situ*^26,27^ as well as after islet transplantation to the anterior chamber of the eye^28,29^. These studies have confirmed functional similarities between *in vivo* and *in vitro* islets and that glucose-stimulated beta cells within the same islet synchronize^18,27^. However, they also highlight a notable difference that islets exhibit activity at lower glucose concentrations *in vivo* than *in vitro*^18,27^, which likely reflects the presence of other beta cell stimulators than glucose in the *in vivo* situation. We now focused on exploring the population dynamics of islets in vivo. To gain a more comprehensive understanding of the activity of different cell types in multiple islets in a living mouse, it was essential to integrate single-cell resolution methods with large-field imaging.

In vivo Ca²⁺ studies of diabetic islets have been promoted by the availability of transgenic diabetic mouse models expressing Ca^2+^ biosensors in islet cells. In vitro studies have shown that disruptions glucose-induced Ca^2+^ rhythmicity correlates with diabetes pathogenesis^30^. Using *ob/ob* mice expressing the Ca^2+^ biosensor *GCaMP6f* in the beta cells we found that they showed slow Ca^2+^ oscillations which failed to transform into fast ones when the insulin infusion normalized blood glucose from the characteristic hyperglycemia. Under *in vivo* conditions the β-cells in *ob/ob* mouse islets also had a lower glucose activation threshold with Ca^2+^ oscillations at 3-5 mM of sugar.

Our study shows that the islet rhythmicity and their synchrony across pancreas play critical and central roles in precisely maintaining the stability of blood glucose.

## Acknowledge

We are grateful to Erik Gylfe for discussions and critical reading of the manuscript. We thank Anders Tengholm and Beichen Xie for helpful discussions. This work was supported by funding from Chinese Institutes for Medical Research, Beijing to H.R. and the National Natural Science Foundation of China (12090053, 32088101, and T2322026). This work was supported by Talent Plan of Shanghai Branch, Chinese Academy of Sciences (CASSHB-QNPD-2023-026). The authors acknowledge the support from the Spectrometry Core Facility and Shared Instrumentation Core Facility at the Chinese Institutes for Medical Research and the Hangzhou Institute of Medicine (HIM), Chinese Academy of Sciences.

## Author Contributions Statement

H.R. conceived and supervised the study, and wrote the manuscript.

Y.D. and Z.F. designed and performed in vivo experiments, carried out data analysis and wrote the manuscript.

X.W., X.W. and Y.Q. designed and performed slice experiments and carried out data analysis.

S.Y. and W.H. designed and performed isolated islet experiments.

L.H. and performed CGM experiments.

W.Q. contributed to in vivo experiments supervision.

Y.D., L.C., C.T. and X.P. contributed to Figure preparation and experiment supervision.

## Methods

### Animals

*Ins1-Cre^+/-^;GCaMP6f^f/+^* mice (aged 2–5 months) were generated by crossing *Ins1-Cre* mice (Jackson Laboratory, Strain#: 026801) and *Rosa26-GCaMP6f*^flox^ mice (Jackson Laboratory, Strain#: 028865) lines. *Ins1-Cre^+/-^;GCaMP6f^f/+^;ob/ob* mice were crossed by *Ins1-Cre* mice, *Rosa26-GCaMP6f*^flox^ mice and *B6/JGpt-Lep^em1Cd^*^25^*/Gpt* mice (*B6-ob* mice, purchased from GemPharmatech, Nanjing, China). And their heterozygous littermates (*Ins1-Cre^+/-^;GCaMP6f^f/+^;ob/wt*) were used as controls. *Ins1-Cre^+/-^;GCaMP6f^f/+^;*Gcgr^f/f^ mice were crossed by *Ins1-Cre* mice, *Rosa26-GCaMP6f*^flox^ mice and Gcgr knockout mice^21^. The animals were maintained in a specific pathogen-free animal facility at Capital Medical University, which was housed in a 12-hour light/12-hour dark cycle with a temperature of 22°C and humidity levels of 40-60%. The animals had free access to water and chow diet. The experiments were approved by the Ethics Committee of Capital Medical University and performed at the animal facility of the same institution, which had accreditation from the Association for Assessment and Accreditation of Laboratory Animal Care International.

### Continuous Blood Glucose Monitoring

Intra- and inter-day blood glucose fluctuations in mice were monitored by continuous glucose monitors (CGMs, Kangtai, GX-01S) immobilized on the back of the mice. CGMs were fitted to both wild-type and *ob/ob* mice used in the above experiments. After anesthetizing the mice, the hair on their backs was shaved. A small incision was made in the skin on one side of the mouse’s back, and the epidermis was bluntly separated from the subcutaneous tissue using forceps. The CGM probe was implanted subcutaneously, and the monitor was sutured and fixed on the mouse’s back with medical non-absorbable sutures. On the first and second days after surgery, meloxicam was administered via intraperitoneal injection for anti-inflammatory and analgesic purposes. The CGM was calibrated using tail vein blood glucose on the day following surgery. After calibration, blood glucose was continuously monitored at a rate of one measurement per min for 8-15 days with a under conditions of free movement and ad Libitum Feeding for each mouse.

### Jugular Vein Cannulation

Glucose and insulin administered during in vivo imaging were delivered via jugular vein cannulation, surgical procedures on mice are required prior to imaging. After anesthetizing the mice, the hair on the mouse neck was shaved, and an incision was made to expose one side of the jugular vein. A 27G indwelling needle filled with 0.2% heparinized saline was inserted into the unilateral jugular vein. A small amount of heparinized saline was flushed to clear backflow of blood from the needle lumen, followed by clamping the stopcock. Syringes containing glucose solutions and insulin were mounted on a microflow pump, connected to the tail end of the indwelling needle via a needle-tipped microflow tube to precisely control drug delivery during imaging experiments. Injection volume and rate were regulated by pre-programmed MATLAB protocols.

### In vivo Multi-islet Calcium Fluorescence Imaging

We performed in vivo multi-islet calcium fluorescence imaging in *Ins-cre^+/-^; GCaMP6f^f/+^* mice (C57BL/6 background), enabling simultaneous observation of 20-100 islets for >5 h. After a 16 h overnight fast, mice were anesthetized with 1.25% tribromoethanol and placed on a heating pad, an incision was made in the middle of upper abdomen. The pancreas exteriorized and placed underneath a height-adjustable microscopy slide and the position is adjusted to hold the pancreas in place. The pancreas was imaged on the Mshot MZX81 microscope using 2× objective, and capturing using a Mshot MS23 camera. We capture one image per second. We acquire ∼2 h of baseline imaging before administering an intravenous glucose bolus (0.6 g/kg) to test the glucose tolerance (IVGTT). Two hours post-IVGTT, we infused 1.1M glucose at escalating rates (30→40→50 mg/kg/min, 20 min each). Blood glucose was monitored via tail vein measurements. Real-time pancreatic islet beta-cells Intracellular calcium concentration measured as GCaMP6 fluorescence intensity in each ROI.

### In vivo Multi-islet Calcium Fluorescence Imaging in *ob/ob* mice

Jugular vein cannulation and surgical procedures were performed on ob/ob mice as previously described. During baseline imaging, to match the blood glucose levels of *ob/ob* and WT mice, insulin (0.2 U/kg) was intravenously infused at 30 minutes after the start of imaging, with the total dose administered over 10 minutes. After waiting for blood glucose levels to drop to 5–10 mM, an intravenous glucose tolerance test (IVGTT, 0.6 g/kg) was conducted. Following the return of blood glucose to pre-IVGTT levels, 1.1M glucose was infused at escalating rates (30→40→50 mg/kg/min, 20 minutes each).

### Micro-glucose Pulse Test

Under imaging operations as described above, when the calcium oscillation pattern of most islets shifted to a fast condition, micro-glucose pulse (20-40 mg/kg) was given intravenous approximately every 30 min, each pulse was administered over 15 seconds.

### Pancreatic slice Calcium imaging

Low-gelling agarose (0.3 g in 20 mL extracellular solution), was melted and kept at 37°C. The animals were anesthetized by intraperitoneal injection of tribromoethanol, the abdominal cavity opened and 2.5 ml agarose injected into the distally clamped bile duct. After injection, the pancreas was cooled with an ice-cold extracellular solution. The extracellular solution consisted of (mM): 125 NaCl, 2.5 KCl, 26 NaHCO3, 1.25 NaH2PO4, 2 Na-pyruvate, 0.5 ascorbic acid, 3 myo-inositol, 6 lactic acid, 1 MgCl2, and 2 CaCl2. The tissue was transferred to a dish filled with agarose and immediately cooled on ice. The agarose-embedded pancreatic tissue was then glued onto the sample plate of the vibrotome (VT 1000 S, Leica, Nussloch, Germany). 270 μm thick slices were cut using a vibratome (VT1200S, Leica) at 85 Hz. The cut speed is 0.08 mm/sec. During slicing the tissue was kept in an ice-cold extracellular solution bubbled with a gas mixture containing 95% O2 and 5% CO2 at barometric pressure to ensure oxygenation and a pH of 7.4. After slicing the tissue slices were used for imaging.

Imaging was performed on Olympus SpinSR10 Xplore (10x, NA 1.0). GCaMP6f was excited by an argon 488 nm laser. 16-bit 2304×2304 pixels images were acquired every 3 seconds. Large-field imaging was performed using a Mshot MZX81 microscope with a 2x objective lens and photographed using a Mshot MS23 camera. This stimulation is Krebs-Ringer buffer (KRB) solution (125 mM NaCl, 5.9 mM KCl, 2.56 mM CaCl2*2H20, 1.2 mM MgCl2*6H20, 25 mM HEPES, 1 mM Glutamine, pH7.2-7.4) with glucose of 1 mM (15 min), 3 mM (15 min), 5 mM (15 min), 7 mM (15 min), 10 mM (30 min), 20 mM (30 min) concentrations of glucose, 25 mM KCl (15 min) and 150 uM diazoxide (15 min). The solution was gassed with a gas mixture containing 95% O_2_ and 5% CO_2_ at a flow rate of 3.3 ml/min in 50 mL solution, and the solution temperature was maintained at 37°C.

### Isolated islet Calcium imaging

The mice were euthanized by cervical dislocation, and following the instillation of 2 ml of 0.25% trypsin from the common pancreatic duct, the entire pancreas was extracted from the mouse. Primary islets were isolated by means of digestion at 37°C for a period of fifteen minutes, after which they were washed in KRB solution (125 mM NaCl, 5.9 mM KCl, 2.56 mM CaCl2*2H20, 1.2 mM MgCl2*6H20, 25 mM HEPES, 1 mg/ml BSA and 1 mM Glutamine, pH7.2-7.4). The islets were cultured at a temperature of 37°C in RPMI 1640 medium (5% CO_2_), containing 10 mmol/L glucose, 10% fetal bovine serum, 100 μg/mL streptomycin, and 100 IU/mL penicillin.

Pancreatic islets were injected into the microfluidic chip, imaged using a Nikon ECLIPSE Ji with a 475-nm laser, an intensity of 20%, and an exposure time of 100 milliseconds. The islets were maintained at 37°C, 95% O_2_ and 5% CO_2_ during imaging. The reagents were then introduced into the microfluidic chip by means of a TS-1B syringe pump (LongerPump), at a flow rate of 200 uL/h. This process was controlled by MATLAB software.

### Semaglutide treatment

Semaglutide treatment of the mice was performed by daily i.p. injections of 0.3 mg/kg body weight (AdipoGen, cat. 910463-68-2) for 1 week, and sham treatment was performed by i.p. injection of NaCl.

**Video. 1** In vivo imaging of *Ins1^+/-^;GCaMP6f^f/+^* mice.

Hyperglycemia-to-Euglycemia Induces a coordinated Slow-to-Fast (HESF) Islet Ca^2+^ Oscillation Transition In Vivo. Blood glucose, 3 islet Ca^2+^ traces from *Ins1^+/-^;GCaMP6f^f/+^* mice are shown.

## References

1. Lehrstrand, J. et al. Illuminating the complete ß-cell mass of the human pancreas-signifying a new view on the islets of Langerhans. Nat. Commun. 15, 3318 (2024).

2. Zhu, L., et al. Intraislet glucagon signaling is critical for maintaining glucose homeostasis. JCI Insight 4, e127994 (2019).

3. Tariq, M. et al. Prolonged culture of human pancreatic islets under glucotoxic conditions changes their acute beta cell calcium and insulin secretion glucose response curves from sigmoid to bell-shaped. Diabetologia 66, 709–723 (2023).

4. Pedersen, M. G., Bertram, R. & Sherman, A. Intra- and Inter-Islet Synchronization of Metabolically Driven Insulin Secretion. Biophys. J. 89, 107– 119 (2005).

5. Carroll, P. B., Zeng, Y., Alejandro, R., Starzl, T. E. & Ricordi, C. Glucose homeostasis is regulated by donor islets in xenografts. Transplant. Proc. 24, 2980–2981 (1992).

6. Rodriguez-Diaz, R. et al. Paracrine Interactions within the Pancreatic Islet Determine the Glycemic Set Point. Cell Metab. 27, 549–558.e4 (2018).

7. Huang, J. L. et al. Paracrine signalling by pancreatic δ cells determines the glycaemic set point in mice. Nat. Metab. 6, 61–77 (2024).

8. Ashcroft, F. M., Harrison, D. E. & Ashcroft, S. J. H. Glucose induces closure of single potassium channels in isolated rat pancreatic β-cells. Nature 312, 446– 448 (1984).

9. Cook, D. L. & Hales, N. Intracellular ATP directly blocks K+ channels in pancreatic B-cells. Nature 311, 271–273 (1984).

10. Rorsman, P. & Braun, M. Regulation of insulin secretion in human pancreatic islets. Annu. Rev. Physiol. 75, 155–179 (2013).

11. Nunemaker, C. S. et al. Glucose Modulates [Ca2+]i Oscillations in Pancreatic Islets via Ionic and Glycolytic Mechanisms. Biophys. J. 91, 2082–2096 (2006).

12. Gylfe, E. & Tengholm, A. Neurotransmitter control of islet hormone pulsatility. Diabetes Obes. Metab. 16, 102–110 (2014).

13. Berggren, P.-O. et al. Removal of Ca2+ Channel β3 Subunit Enhances Ca2+ Oscillation Frequency and Insulin Exocytosis. Cell 119, 273–284 (2004).

14. Gilon, P. The Role of α-Cells in Islet Function and Glucose Homeostasis in Health and Type 2 Diabetes. J. Mol. Biol. 432, 1367–1394 (2020).

15. Dai, X.-Q. et al. Heterogenous impairment of α cell function in type 2 diabetes is linked to cell maturation state. Cell Metab. 34, 256–268.e5 (2022).

16. Rorsman, P. & Ashcroft, F. M. Pancreatic β-Cell Electrical Activity and Insulin Secretion: Of Mice and Men. Physiol. Rev. 98, 117–214 (2018).

17. Hellman, B., Dansk, H. & Grapengiesser, E. Somatostatin promotes glucose generation of Ca2+oscillations in pancreatic islets both in the absence and presence of tolbutamide. Cell Calcium 74, 35–42 (2018).

18. Jacob, S. et al. In vivo Ca^2+^ dynamics in single pancreatic β cells. FASEB J. 34, 945–959 (2020).

19. Capozzi, M. E., et al. β cell tone is defined by proglucagon peptides through cAMP signaling. JCI Insight 4, e126742 (2019).

20. Svendsen, B. et al. Insulin secretion depends on intra-islet glucagon signaling. Cell Rep. 25, 1127–1134.e2 (2018).

21. Zhang, Y. et al. Glucagon potentiates insulin secretion via β-cell GCGR at physiological concentrations of glucose. Cells 10, 2495 (2021).

22. Ren, H. et al. Pancreatic α and β cells are globally phase-locked. Nat. Commun. 13, 3721 (2022).

23. Liu, Y.-J., Tengholm, A., Grapengiesser, E., Hellman, B. & Gylfe, E. Origin of slow and fast oscillations of Ca2+ in mouse pancreatic islets. J. Physiol. 508, 471–481 (1998).

24. Dickerson, M. T. et al. Gi/o protein-coupled receptor inhibition of beta-cell electrical excitability and insulin secretion depends on Na+/K+ ATPase activation. Nat. Commun. 13, 6461 (2022).

25. Sánchez-Andrés, J. V., Gomis, A. & Valdeolmillos, M. The electrical activity of mouse pancreatic beta-cells recorded in vivo shows glucose-dependent oscillations. J. Physiol. 486, 223–228 (1995).

26. Ahl, D. et al. Turning Up the Heat: Local Temperature Control During in vivo Imaging of Immune Cells. Front. Immunol. 10, 2036 (2019).

27. Fernandez, J. & Valdeolmillos, M. Synchronous glucose-dependent [Ca ^2+^]_i_ oscillations in mouse pancreatic islets of langerhans recorded in vivo. FEBS Lett. 477, 33–36 (2000).

28. Ilegems, E. et al. Reporter islets in the eye reveal the plasticity of the endocrine pancreas. Proc. Natl. Acad. Sci. 110, 20581–20586 (2013).

29. Speier, S. et al. Noninvasive in vivo imaging of pancreatic islet cell biology. Nat. Med. 14, 574–578 (2008).

30. Corbin, K. L., Waters, C. D., Shaffer, B. K., Verrilli, G. M. & Nunemaker, C. S. Islet Hypersensitivity to Glucose Is Associated With Disrupted Oscillations and Increased Impact of Proinflammatory Cytokines in Islets From Diabetes-Prone Male Mice. Endocrinology 157, 1826–1838 (2016).

